# Evaluation of a novel multiplexed assay for determining IgG levels and functional activity to SARS-CoV-2

**DOI:** 10.1101/2020.07.20.213249

**Authors:** Marina Johnson, Helen R. Wagstaffe, Kimberly C. Gilmour, Annabelle Lea Mai, Joanna Lewis, Adam Hunt, Jake Sirr, Christopher Bengt, Louis Grandjean, David Goldblatt

**Affiliations:** Great Ormond Street Institute of Child Health, University College London, 30 Guilford Street, London, WC1N 1 EH, UK; Great Ormond Street Children’s Hospital NHS Foundation Trust, Great Ormond Street, London, WC1N 3JH, UK

## Abstract

**Background:** The emergence of SARS-CoV-2 has led to the development of new serological assays that could aid in diagnosis and evaluation of seroprevalence to inform an understanding of the burden of COVID-19 disease. Many available tests lack rigorous evaluation and therefore results may be misleading.

**Objectives:** The aim of this study was to assess the performance of a novel multiplexed immunoassay for the simultaneous detection of antibodies against SARS-CoV-2 trimeric spike (S), spike receptor binding domain (RBD), spike N terminal domain and nucleocapsid antigen and a novel pseudo-neutralisation assay.

**Methods:** A multiplexed solid-phase chemiluminescence assay (Meso Scale Discovery) was evaluated for the simultaneous detection of IgG binding to four SARS-CoV-2 antigens and the quantification of antibody-induced ACE-2 binding inhibition (pseudo-neutralisation assay). Sensitivity was evaluated with a total of 196 COVID-19 serum samples (169 confirmed PCR positive and 27 anti-nucleocapsid IgG positive) from individuals with mild symptomatic or asymptomatic disease. Specificity was evaluated with 194 control serum samples collected from adults prior to December 2019.

**Results:** The specificity and sensitivity of the binding IgG assay was highest for S protein with a specificity of 97.4% and sensitivity of 96.2% for samples taken 14 days and 97.9% for samples taken 21 days following the onset of symptoms. IgG concentration to S and RBD correlated strongly with percentage inhibition measured by the pseudo-neutralisation assay.

**Conclusion:** Excellent sensitivity for IgG detection was obtained over 14 days since onset of symptoms for three SARS-CoV-2 antigens (S, RBD and N) in this multiplexed assay which can also measure antibody functionality.

## Introduction

Severe acute respiratory syndrome-related coronavirus-2 (SARS-CoV-2) was first recognised in January 2020 and rapidly spread world-wide with the WHO declaring a COVID-19 pandemic on March 11^th^, 2020 (1). Soon after the identification and genetic sequencing of the virus, diagnostic tests became available for the detection of live virus in human secretions followed rapidly by tests designed to measure antibodies to SARS-CoV-2 antigens. Antibody tests have a variety of uses including supporting diagnosis and informing individual risk of future disease and thereby determining correlates of and duration of protection. With further potential for understanding exposure to virus which in turn could help inform disease burden estimates, studies of transmission dynamics and modelling of the epidemic. Antibody tests are particularly important in the context of mild or asymptomatic disease where a swab reverse transcriptase polymerase chain reaction (RT-PCR) test may be negative. For this reason, an understanding of the sensitivity and specificity of the tests being used is critical.

The trimeric spike (S) protein of SARS-CoV-2 is a large molecule that is critical to virus dissemination and pathogenesis. It is densely glycosylated and present on the viral surface and in most cases is cleaved by host proteases into the S1 and S2 subunits, which are responsible for receptor recognition and membrane fusion respectively. S1 uses a region of the molecule, known as the receptor binding domain (RBD) to bind to host ACE-2 receptor and thereby gain entry to the cell (2). Due to this critical function in host-cell entry, the S protein is a major target for vaccine research. The N terminal domain (NTD) of the spike protein does not interact with the receptor but contains the functional elements required for membrane fusion of the virion. The nucleocapsid (N) protein plays an important role in transcription enhancement and viral assembly (3).

Specific immunoglobulin-G (IgG) and IgM antibody responses to SARS-CoV-2 S, N and RBD of the spike protein develop between 6-15 days following disease-onset (4). Despite a rapid increase in the number and availability of serologic assays that can detect antibodies against SARS-CoV-2, most have undergone minimal external evaluation and validation (5). The high sensitivity and specificity for commercially obtainable kits are often not reproduced when appropriate samples are used for evaluation. A recent large scale Spanish seroprevalence study used a point of care IgG test with a stated sensitivity of 97.2% but on verification found it to have a sensitivity of either 82.1%, 89.7%, 99.6% or 100% depending on the sample sets used for evaluation (6). All assays currently suffer from the absence of a defined standard serum so results are reported as positive or negative or as optical density readouts complicating the comparison between assays and studies. Furthermore, most assays measure responses to a single antigen, usually nucleocapsid or spike/spike derived proteins, which may not capture the breadth of antibody responses to SARS-CoV-2. Finally, for many binding assays, the relationship between the concentration of antibody detected and their function is unclear and few available assays permit the measurement of both binding and function on the same testing platform.

We have evaluated a novel assay designed to simultaneously measure IgG to four SARS-CoV-2 antigens; full-length trimeric S, RBD and NTD of spike as well as N protein. The assay, based on Meso Scale Discovery (MSD) technology, utilises a 96-well based solid-phase antigen printed plate and an electrochemiluminescent detection system. In addition, unlike most binding assays, this assay can be adapted to measure the ability of serum to inhibit the interaction between spike protein components and soluble ACE-2, also called a pseudo-neutralisation assay (7). To evaluate the sensitivity and specificity of the MSD assay, we were able to utilise a relatively large number of samples obtained from SARS-CoV-2 RT-PCR positive health care workers or patients as well as antibody positive health care staff enrolling in a large SARS-CoV-2 cohort study.

## Materials and Methods

### Serum Samples

Serum samples for sensitivity analyses were obtained from Great Ormond Street Children’s Hospital NHS Foundation Trust (GOSH) and came from three sources; (i) healthcare workers who tested SARS-CoV-2 RT-PCR positive following signs or symptoms of COVID-19 and who gave written consent for participation in the service evaluation of SARS-CoV-2 serological assays, (ii) staff enrolling in a prospective longitudinal cohort study of SARS-CoV-Serology (COSTARS, IRAS 282713, ClinicalTrials.gov Identifier: NCT04380896) who tested positive in a commercial screening assay for anti-Nucleocapsid IgG (Epitope Diagnostics Inc, San Diego, USA) (iii) a small number of RT-PCR positive sera from hospitalised children (n=10).

Serum samples for the analysis of specificity were collected prior to December 2019 and derived from anonymised samples in assay development or quality control sera developed for other assays or residual, anonymised samples from healthy adults enrolled in previous studies.

Serum from two individuals with high convalescent antibody levels were pooled to create an interim standard serum. This serum was calibrated against research reagents NIBSC 20/130 and NIBSC 20/124 distributed by the National Institute for Standards and Biological Control (NIBSC, Potters Bar, UK, https://www.nibsc.org/) for the purpose of development and evaluation of serological assays for the detection of antibodies against SARS-CoV-2. These two plasma samples were obtained from COVID-19 recovered patients and were distributed with known end-point titres to trimeric S, S1 and N as well as antibody functionality measured by live virus neutralisation, pseudo-neutralisation and plaque reduction neutralisation.

### Serological assays

Samples were screened for IgG to SARS-CoV-2 N protein using a commercially available kit (Epitope Diagnostics Inc, San Diego, USA) as previously described (8).

### Meso Scale Discovery coronavirus panel for COVID-19 serology

A multiplexed MSD immunoassay (MSD, Rockville, MD) was used to measure the antigen-specific response to SARS-CoV-2 infection and other respiratory pathogens. A MULTI-SPOT® 96-well, 10 Spot Plate was coated with four SARS CoV-2 antigens (S, RBD, NTD and N), SARS-CoV-1 and MERS spike trimers, spike proteins from seasonal coronaviruses OCV43S and HKU1, influenza A antigen derived from H3/HongKong and Bovine Serum Antigen. Antigens were spotted at 200-400 µg/mL in a proprietary buffer, washed, dried and packaged for further use (MSD® Coronavirus Plate 1). Proteins were expressed in a mammalian cell expression system (Expi 293F), purified by ion exchange chromatography, affinity purification, and size exclusion chromatography. They were characterized by reducing SDS Page chromatography, mass spectrometry, size-exclusion chromatography and multi-angle light scattering (SEC-MALS). All protein constructs were produced with His6 and/or Strept-TAG affinity tags to support affinity purification; the spike proteins were produced as trimers in the pre-fusion form. These assays were developed by MSD in collaboration with the Vaccine Research Center at NIAID (A. McDermott).

Internal quality controls and reference standard reagents were developed from pooled human serum. To measure IgG antibodies, plates were blocked with MSD Blocker A for between 30 minutes and 2 hours then washed three times prior to the addition of reference standard, controls and samples diluted 1:500 in diluent buffer. After incubation for 2 hours with shaking at 700rpm, the plates were washed three times and detection antibody was added at 2 µg/mL (MSD SULFO-TAG™ Anti-Human IgG Antibody). Plates were incubated for 1 hour with shaking and washed three times. MSD GOLD™ Read Buffer B was added and the plates were read using a MESO® SECTOR S 600 Reader.

### Meso Scale Discovery pseudo-neutralisation assay

Plates were blocked and washed as above, assay calibrator (COVID-19 neutralising antibody; monoclonal antibody against S protein; 200µg/ml), control sera and test sera samples diluted 1 in 10 in assay diluent were added to the plates. Plates were incubated for 1 hour with shaking at 700rpm. A 0.25µg/ml solution of MSD SULFO-TAG™ conjugated ACE-2 was added to unwashed plates followed by incubation for 1 hour with shaking, plates were washed and read as above. Percentage inhibition was calculated relative to the assay calibrator; the maximum inhibition reached with calibrator was set as 100% inhibition, minimum at 0.01%.

### Statistical analysis

Statistical analysis was performed using MSD Discovery Workbench and GraphPad Prism version 8.0 (GraphPad, San Diego, CA). Antibody concentration in arbitrary units (AU) was interpolated from the ECL signal of the internal standard sample using a 4-parameter logistic curve fit. ROC curves showing the sensitivity and specificity (plotted as 100%-specificity %) calculated using each value in the data as a cut-off were plotted for each antigen. A cut-off antibody concentration was chosen based on the lowest value leading to a positive likelihood ratio (LR) of >10, in order to maximise sensitivity while providing strong evidence to rule-in infection (9). For S antigen binding, all LR’s were above 10, therefore the LLOD was used as the cut-off for this antigen. Positive predictive value (PPV) was calculated as

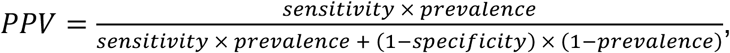

negative predictive value (NPV) was calculated as

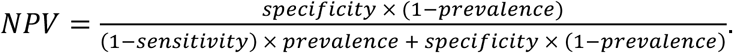

Comparisons between groups were performed by Kruskal-Wallis one-way ANOVA with Dunn’s correction for multiple comparisons. Correlation analysis was performed using Spearman correlation. P values of <0.05 were considered as significant. Latent class models with two classes were fitted with the binary antibody responses as outcome variables, using the poLCA package in the R statistical environment. The code used for the latent class analysis is available on request.

## Results

### Participants and samples

SARS-CoV-2 positive samples (COVID-19 cohort) comprised 169 PCR positive and 27 anti-N IgG positive serum samples from mild symptomatic or asymptomatic cases (total n=196). The cohort comprised of 138 females, 56 males (2 not recorded) with a median age of 37 years (range 1-66y). Recorded symptoms included abnormal taste and smell, cough, fatigue and fever. The date of symptom onset was established and verified for 168 subjects, time between symptom onset and sampling ranged from 4 to 63 days. Of the 169 individuals with documented RT-PCR testing, 37 samples were negative for nucleocapsid IgG on the EDI screening ELISA and 11 were equivocal. Serum samples were collected between 26^th^ March and 18^th^ May 2020 and analysed between 1^st^ June and 10^th^ July 2020.

Control serum samples for the analysis of specificity comprised 194 anonymised legacy samples obtained from healthy adults, aged predominantly over 50 years.

### Standard serum assignment

An internal standard serum was assigned values for S, RBD and N by calibration against the NIBSC control sera. The ECL signal obtained for NIBSC 20/130 was used as a binding curve to assign arbitrary unit (AU) values for S and RBD while NIBSC 20/124 was used to assign a value for N (Supplementary Figure S1). Binding of pooled standard serum to NTD produced low ECL signals and no endpoint titre corresponding to NTD antigen was available for standard serum assignment. The interim values assigned were S 2154 AU, RBD 1837 AU and N 3549 AU. NTD and the remaining antigens were assigned a value of 1000 AU. The focus of this study was the evaluation of the four SARS-CoV-2 antigens only.

### Evaluation of the coronavirus panel for COVID-19 serology

The lower limit of detection (LLOD) was assigned as 1% of the standard value in AU, for statistical purposes, values below LLOD were reported as half LLOD (Table 1). The upper limit of detection (ULOD) was assigned for NTD and RBD only as the S and N antigen did not reach an upper limit (Table 1). For statistical purposes, ULOD was assigned the highest calculated concentration plus 20%.

**Table 1:** The lower limit of detection (LLOD), upper limit of detection (ULOD), quality control (QC) sample range in arbitrary units (AU) and positive/negative cut-off for each SARS-CoV-2 antigen analysed.

| Antigen | LLOD (max.)<br>(AU) | ULOD (min.)<br>(AU) | QC sample<br>range (AU) | Positive/<br>negative cut-off |
| --- | --- | --- | --- | --- |
| CoV-2 S | 21.54 | NA | 1092-1478 | 21.5 |
| CoV-2 RBD | 18.37 | 125477 | 2176-2944 | 201.7 |
| CoV-2 N | 35.49 | NA | 3627-4907 | 185.4 |
| CoV-2 NTD | 10.00 | 19452 | 1004-1359 | 1924 |

The coefficient of variation (CV) between duplicates was assessed by analysing 390 samples run on 11 plates on 3 different days. All antigens produced a mean CV of <15%, with only NTD falling above the accepted CV of 15% at 17.4% (data not shown). Intra-assay (within plate) and inter-assay (between plate) variation of the assay was assessed by running four samples of varying antibody levels in four replicates on the same plate and across 4 different runs on different days (Supplementary Table 1). The mean intra-assay CV was 6.2% and inter-assay variation <15% across all SARS-CoV-2 antigens except NTD (19.0%) on one of four samples.

To control day to day performance of the assay, a QC sample was run on each plate and an acceptable performance range was set as within 3 SD of the mean. This was determined by running the sample on 8 different plates on 8 different days (average CV 10.3%) (Table 1).

### Assay sensitivity and specificity

Figure 1A-D shows the concentration of IgG to each SARS-CoV-2 antigen in the COVID-19 cohort and the controls. Receiver Operating Characteristic (ROC) curves were plotted to visualise the trade-off between sensitivity and specificity for each antigen (Figure 2A-D). The high area under the curve (AUC) values for S (0.95%; 95%CI 0.93 to 0.97), RBD (0.92%, 0.89-0.95) and N (0.90%, 0.87-0.94) indicates the high accuracy of these tests. Table 1 shows the cut-off values selected using our rule of choosing the lowest cut-off with LR>10. For S all LRs were above 10, therefore the LLOD was used as the cut-off for this antigen. NTD data was less consistent than the other SARS-CoV-2 antigens and demonstrated lower sensitivity and specificity (Figure 2D), so this antigen was not evaluated further.

**Figure 1:**
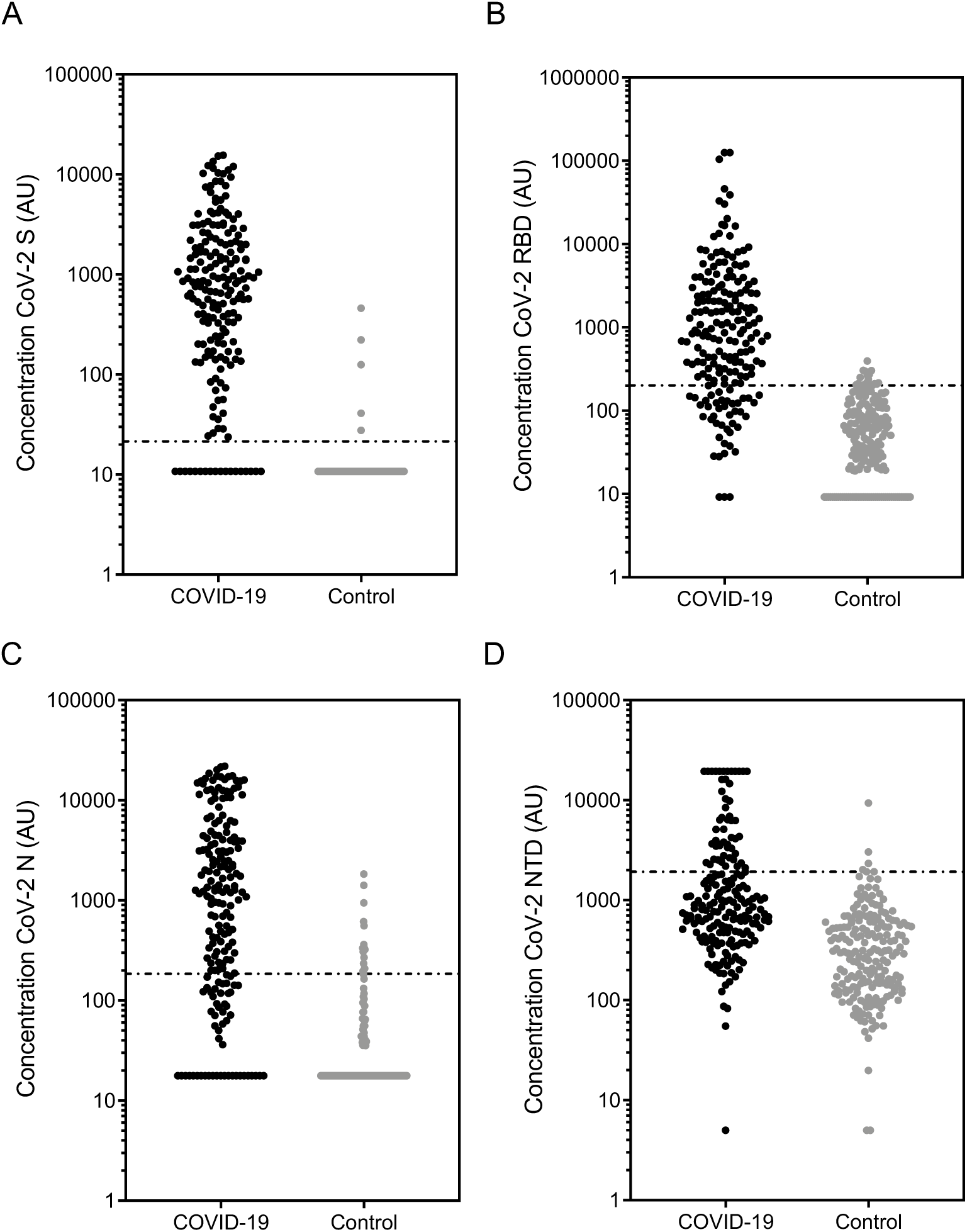
Anti-SARS-CoV-2 IgG concentration. The concentration of SARS-CoV-2 antibody against (a) spike (S), (b) receptor binding domain (RBD), (c) nucleocapsid (N) and (d) N terminal domain (NTD) was measured using the MSD coronavirus panel. Graphs show data in arbitrary units (AU) (based on the calibrated internal standard serum) in the COVID-19 cohort (n=196) and controls (n=194, pre-December 2019). Line shows positive/negative discrimination cut-off.

**Figure 2:**
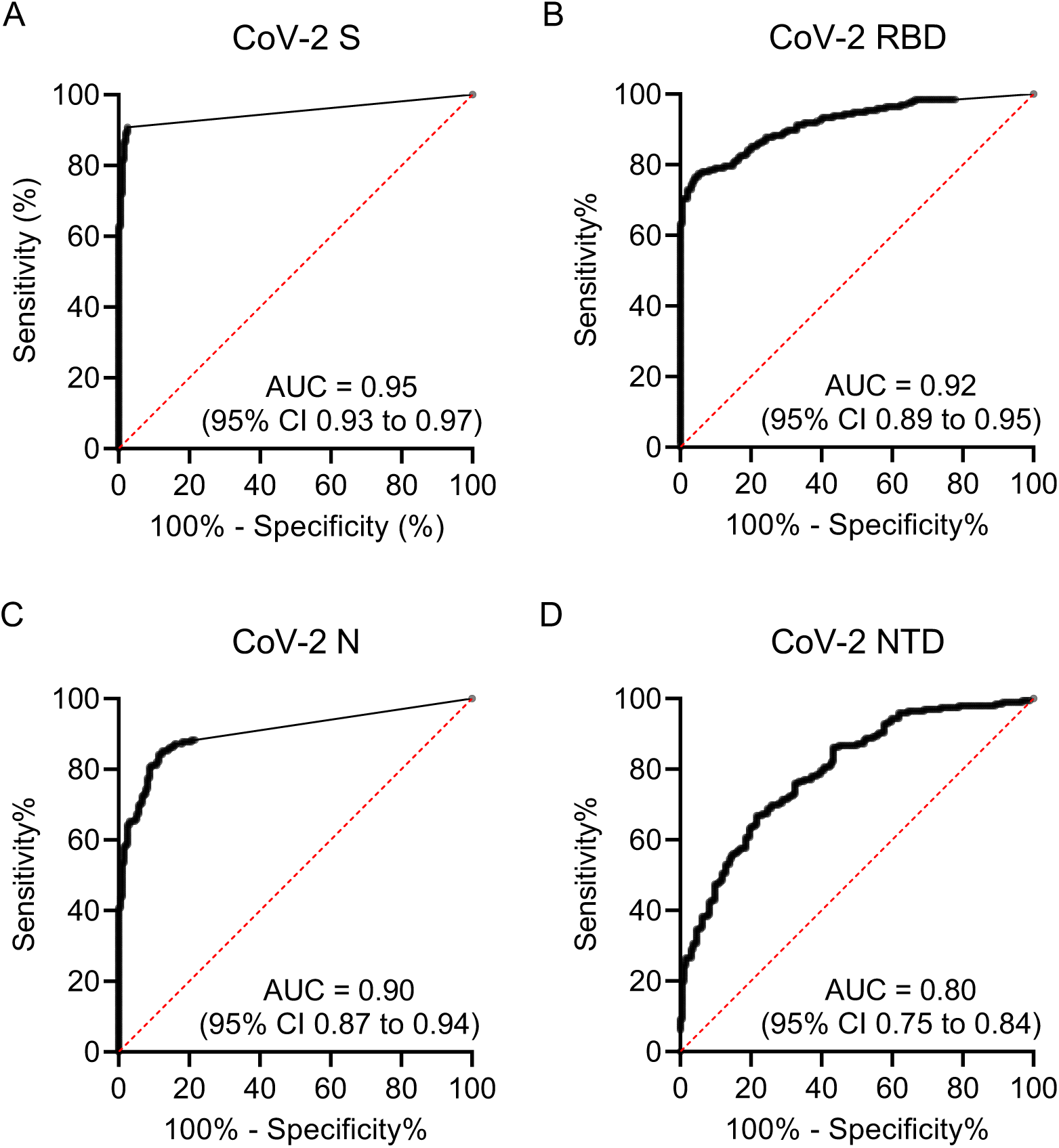
Receiver Operating Characteristic (ROC) curves for each SARS-CoV-2 antigen. Sensitivity and specificity were calculated using each value in the data table as a cut-off value (n=390). Graphs show the sensitivity vs 100%-specificity of SARS-CoV-2 antigen (a) spike (S), (b) receptor binding domain (RBD), (c) nucleocapsid (N) and (d) N terminal domain (NTD). The area under curve (AUC) and 95% CI is also shown for each antigen.

The specificity for S, RBD and N assays calculated from the control sera were 97.4% (95%CI 94.1 to 98.9), 92.3% (95%CI 87.6 to 95.3) and 92.8% (95%CI 88.2 to 95.7) respectively (Table 2). Assay sensitivity was initially calculated on the entire COVID-19 cohort; S antigen had the highest AUC and was the most sensitive and specific at 90.8% and 97.4% respectively.

**Table 2:** Assay specificity calculated for each SARS-CoV-2 antigen from the control cohort.

| Antigen | n | Positive | Negative | Specificity (95% CI)<br>(%) |
| --- | --- | --- | --- | --- |
| CoV-2 S | 194 | 5 | 189 | 97.4% (94.1 to 98.9) |
| CoV-2<br>RBD | 194 | 15 | 179 | 92.3% (87.6 to 95.3) |
| CoV-2 N | 194 | 14 | 180 | 92.8% (88.2 to 95.7) |

Using the calculated specificity and sensitivity, the positive and negative predictive values (PPV and NPV) for each antigen at a range of prevalence estimates between 0.01 and 0.5 were calculated (Supplementary Figure 3A-B). The PPV and NPV were best for S antigen; for an overall prevalence of 10% the assay has a PPV of 80.4% and NPV of 99.6% for samples taken over 14 days since onset of symptoms, this increased to 92.5% and 98.7% for an overall prevalence of 25%.

### Evaluation of sensitivity according to time since onset of symptoms

Figure 3 shows the anti-S, RBD and N IgG concentration split into time since onset of symptom intervals of 0-7 days, 8-14 days, 15-21 days and over 21 days. For all three antigens, the median antibody concentration increased significantly between 8-14 days and over 21 days and all interval groups were significantly (p=<0.0001) higher than the control cohort (Figure 3A-C). There was a significant association between antibody concentration and time since onset of symptoms (SARS-CoV-2 S, Spearman correlation (r)=0.453; SARS-CoV-2 RBD, r=0.478; SARS-CoV-2 N, r=0.392, all p=<0.0001) (Supplementary Figure 2A-C).

**Figure 3:**
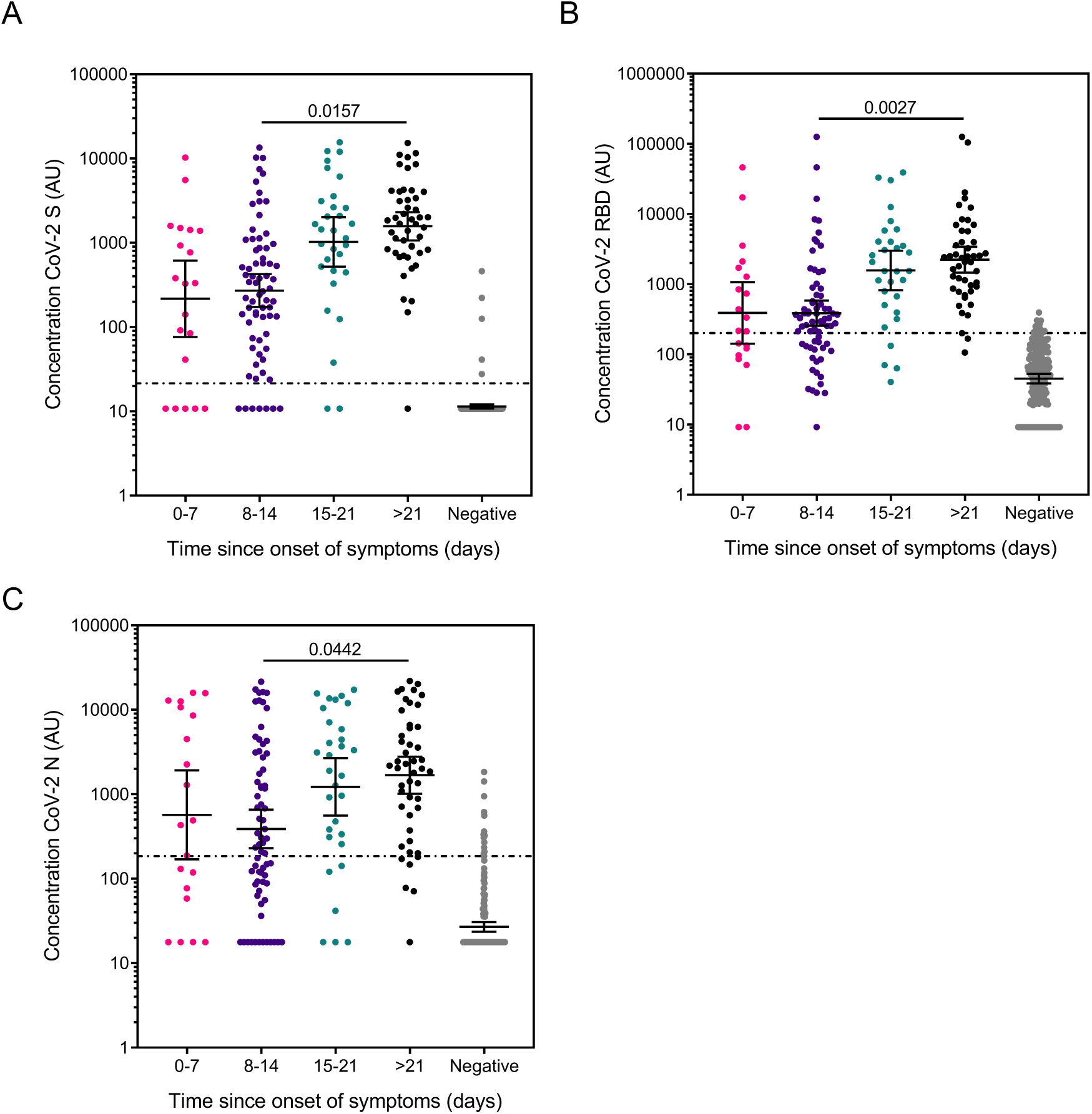
Anti-SARS-CoV-2 IgG concentration according to time since onset of symptoms. Graphs show the concentration of SARS-CoV-2 antibody against (a) spike (S), (b) receptor binding domain (RBD) and (c) nucleocapsid (N) in arbitrary units (AU) (based on the calibrated internal standard serum) of the COVID-19 cohort split in to intervals of 0-7 days, 8-14 days, 15-21 days and over 21 (>21) days since symptom onset (to sample collection). Error bars show geometric mean with 95% CI, line shows positive/negative discrimination cut-off, *p<0.05, ** p<0.01 determined by Dunn’s multiple comparisons test. Comparisons across interval groups had p<0.0001 by one-way ANOVA Kruskal-Wallis test. The assay sensitivity at each time point is shown in Table 3.

The assay cut-off determined above was applied and sensitivity and specificity were calculated for groups 0-7 days, over 7 days, over 14 days and over 21 since the onset of symptom for (Table 2). The S antigen was the most sensitive of the three, with a sensitivity of 96.2% and 97.9% >14 days and >21 days since onset of symptoms respectively.

**Table 3:** Assay sensitivity by time since onset of symptoms for each SARS-CoV-2 antigen calculated using the COVID-19 cohort with verified time between onset of symptoms and blood sampling. Time was divided into 0-7 days, over 7 days, over 14 days and over 21 days since the onset of symptoms.

| Antigen | Group |  | n | Positive | Negative | Sensitivity (95% CI) (%) |
| --- | --- | --- | --- | --- | --- | --- |
| CoV-2 S | Total |  | 196 | 178 | 18 | 90.8% (86.0 to 94.1) |
|  | Time since onset of symptoms | 0-7 days | 20 | 15 | 5 | 75.0% (53.1 to 88.8) |
|  |  | Over 7 days | 148 | 138 | 10 | 93.2% (88.0 to 96.3) |
|  |  | Over 14 days | 78 | 75 | 3 | 96.2% (89.3 to 99.0) |
|  |  | Over 21 days | 47 | 46 | 1 | 97.9% (88.8 to 99.9) |
| CoV-2 RBD | Total |  | 196 | 153 | 43 | 78.1% (71.8 to 83.3) |
|  | Time since onset of symptoms | 0-7 days | 20 | 12 | 8 | 60.0% (38.7 to 78.1) |
|  |  | Over 7 days | 148 | 119 | 29 | 80.4% (73.3 to 86.0) |
|  |  | Over 14 days | 78 | 71 | 7 | 91.0% (82.6 to 95.6) |
|  |  | Over 21 days | 47 | 44 | 3 | 93.6% (82.8 to 97.8) |
| CoV-2 N | Total |  | 196 | 143 | 53 | 73.0% (66.3 to 78.7) |
|  | Time since onset of symptoms | 0-7 days | 20 | 12 | 8 | 60.0% (38.7 to 78.1) |
|  |  | Over 7 days | 148 | 106 | 42 | 71.6% (63.9 to 78.3) |
|  |  | Over 14 days | 78 | 66 | 12 | 84.6% (75.0 to 91.0) |
|  |  | Over 21 days | 47 | 41 | 6 | 87.2% (74.8 to 94.0) |

### Antibody concentration relationship between antigens

The concentration of anti-S, RBD and N antibody all correlated significantly with each other (p<0.0001; Figure 4A-C), the strongest association was between S and RBD (r=0.882) (Figure 4A). Our two-class latent class model built using binary S, RBD and N antigen results predicted known status with 81.1% (95%CI 74.8-86.2) sensitivity and 99.0% (95%CI 95.9-99.8) specificity. It therefore had lower sensitivity and no meaningful improvement in specificity, compared to using the concentration of S antibody alone, with the 21.54 AU cut-off.

**Figure 4:**
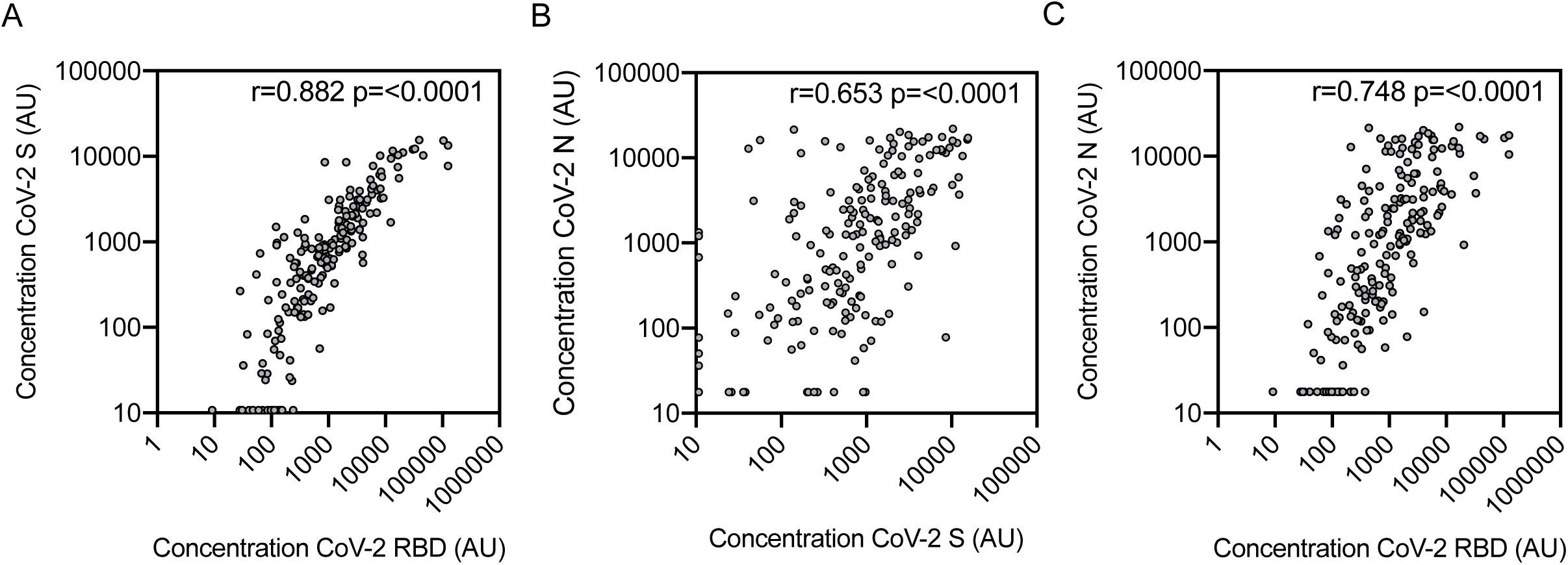
IgG concentration relationship between antigens. Correlation between anti-SARS-CoV-2 antibody concentration of all COVID-19 group samples (n=196) (a) S vs RBD, (b) S vs N and (c) N vs RBD. r and p value were determined by Spearman correlation. p values of <0.05 were considered as significant.

### Pseudo-neutralisation

183 COVID-19 cohort samples with sufficient volume and 194 control group samples were evaluated in the pseudo-neutralisation assay. The percentage inhibition of ACE-2 receptor binding to the S and RBD antigens was calculated for the COVID-19 and control group (Figure 5A-B). The percentage inhibition for the COVID-19 cohort was significantly higher than the controls for both antigens (S, median 1.94% (95%CI 1.36-2.25) vs 0.063% (95%CI 0.053-0.073), p=<0.0001 by Mann-Whitney U test; RBD, 1.50% (95%CI 1.064-2.11) vs 0.38% (95%CI 0.36-0.39); p=<0.0001). In the COVID-19 cohort, there was a significant association between percentage inhibition and IgG concentration for both S and RBD antigens (Spearman correlation (r)=0.805 and r=0.834 respectively, p=<0.0001) (Figure 5C-D).

**Figure 5:**
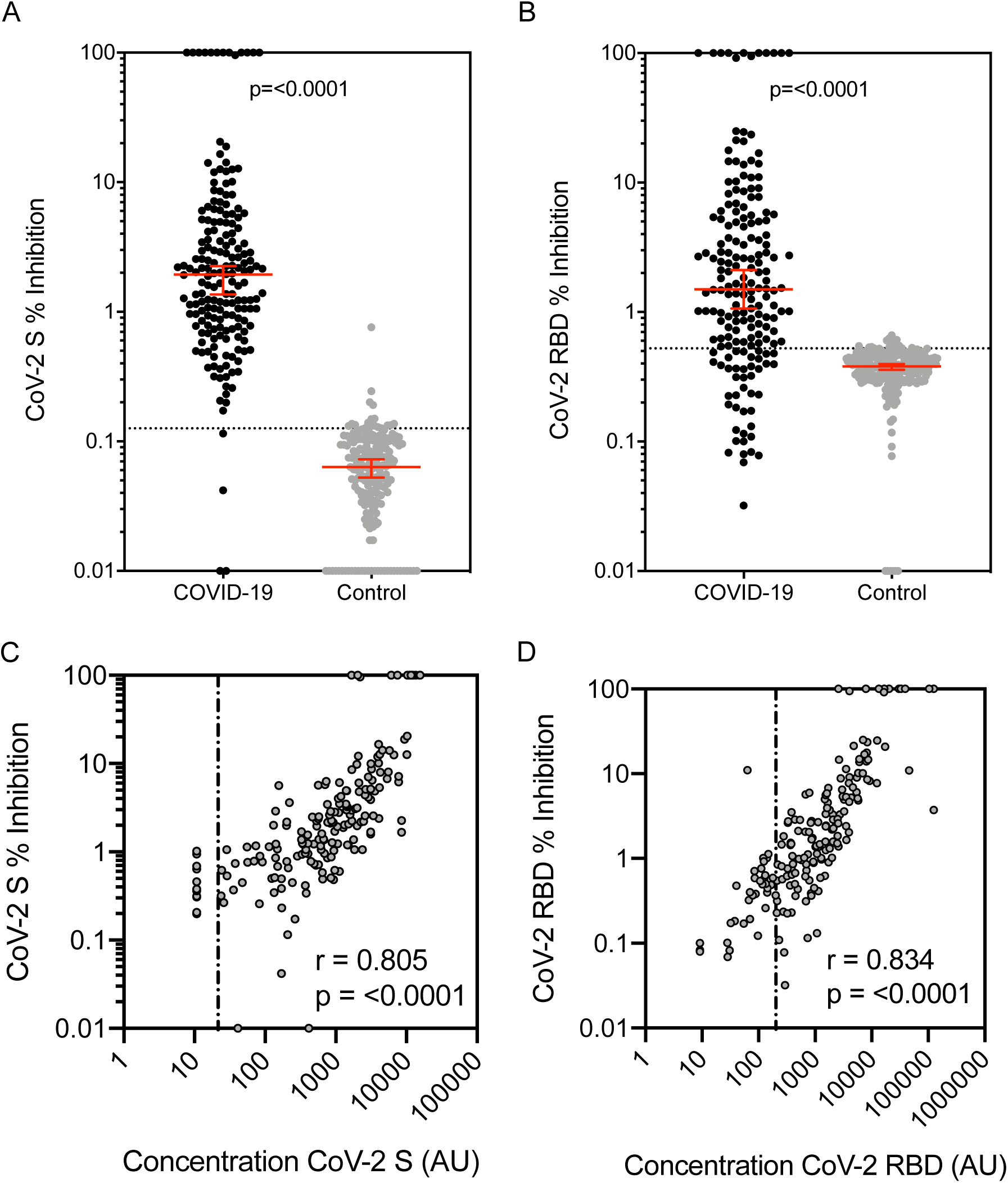
Percentage inhibition by anti-SARS-CoV-2 S and RBD antibody measured by MSD pseudo-neutralisation assay. Inhibition of ACE-2 binding by SARS-CoV-2 antibody against (a) spike (S) and (b) receptor binding domain (RBD) was measured using the MSD coronavirus pseudo-neutralisation assay. 183 COVID-19 cohort samples and 194 control samples were analysed. Graphs show median and 95% CI with a line showing neutralisation assay positive/negative discrimination cut-off determined by ROC. The correlation between antibody concentration and percentage inhibition of (c) S and (d) RBD antigens in all positive group samples was assessed and r and p was determined by Spearman correlation, line shows binding assay positive/negative discrimination cut-off.

ROCs were plotted to visualise the trade-off between sensitivity and specificity for S and RBD neutralisation. Cut-offs (LR>10) were 0.162% for S and 0.524% for RBD (shown by the dotted line on Figure 5A-B). Sensitivity and specificity for S were 97.8% and 97.9% respectively but lower for RBD (77.2% and 92.8% respectively). In the COVID-19 cohort there were some IgG positive sera that did not demonstrate neutralisation (below cut-off, n= 4 for S and 36 for RBD). These sera were predominantly those taken soon after the onset of symptoms; 22 between 0-7 days, 9 over 14 days and 5 over 21 days.

## Discussion

Accurate tests of SARS-CoV-2 antibodies are critical for reliably evaluating exposure to the virus causing COVID-19. Despite a large number of assays rapidly becoming available, many have not undergone rigorous evaluation. In this study we describe a novel assay that can measure antibody to several SARS-CoV-2 antigens simultaneously as well as evaluating the functional capacity of anti-Spike antibodies. The assay we used is based on existing technology developed by Meso Scale Discovery that uses high binding carbon electrodes in the bottom of 96-well microplates. Each well contains up to 10 antigens bound in discrete spots and bound serum-derived IgG is detected by electro-chemiluminescent labelled (SULFO-TAG) anti-human IgG. Electricity is applied to the plate electrodes leading to light emission by the SULFO-TAG labelled detection antibody and light intensity is measured to quantify analytes in the sample. We decided to evaluate IgG only as the kinetics of IgM responses appear to mimic those of IgG and thus add little value (4).

Unlike the majority of studies published to date, we were able to utilise a panel of COVID-19 convalescent plasma recently distributed by WHO to calibrate an internal standard made from pooled convalescent serum. This allowed us to express titres in arbitrary units that can then be compared to other assays that report values calibrated against the WHO panel. The assays performed reliably and consistently over the period of study and passed all the performance criteria expected for a solid-phase based assay with acceptably low inter- and intra-assay coefficients of variation. A QC range established for a medium titre serum gave consistent results throughout the study indicating the stability and repeatability of the platform.

Using a carefully defined cohort of known SARS-CoV-2 exposed individuals and relevant controls we were able to show the sensitivity and specificity of the assay for the four antigens of interest. While all antigens had good specificity, the full-length trimeric spike protein had the highest sensitivity, particularly for serum taken more than 14 days following the onset of symptoms. Comparing our data for the S and RBD antigens to data in a recently published systematic review and metanalysis of the diagnostic accuracy of serological tests for COVID-19 (10) the trimeric spike assay we evaluated had superior sensitivity to all of the assays included in the review while the RBD antigen performance was superior to most. The reason for this could be related to the technical aspects of the assay itself including the integrity of the antigen used and the sensitivity of the detection platform but also the use of a well-defined cohort of individuals with known exposure to SARS-CoV-2. Only one of the four SARS-CoV-2 antigens, the N terminal domain of the spike protein, did not perform well in this assay with poor sensitivity due to the overlap in antibody titres between the COVID-19 cohort and controls.

The ability to simultaneously measure responses to various SARS-CoV-2 antigens could be seen as an advantage in this type of assay although we did not show an advantage of combined analysis of responses to three antigens compared to using S antigen results alone to predict exposure correctly to the virus. The assay format also permitted the measurement of antibody against spike protein derived from SARS-1, MERS and two seasonal coronaviruses, but the results of antibody binding to these antigens could not be assessed in the same way as for the SARS-CoV-2 antigens due to the absence of defined negative and positive serum sets.

A further advantage of this assay is the ability to adapt it for measuring antibody induced inhibition of the interaction between the spike antigen and soluble ACE-2 receptor, without the use of live virus and category 3 facilities. This is important as it is thought to be the major mechanism by which SARS viruses, including SARS-CoV-2 attach to host cell surfaces (11, 12). In the COVID-19 group, there was a good correlation between the concentration of anti-S and anti-RBD IgG and the inhibitory capacity of serum measured in the pseudo-neutralisaton assay, although a few sera bound antigen but did not neutralize ACE-2 binding. Recently, a study of convalescent serum by Sedoux *et al*. identified that the majority of antibodies against spike that were generated during the first weeks of COVID-19 infection were non-neutralising and target epitopes outside the RBD (13) which may account for our results. Few of the control cohort sera had any pseudo-neutralisation activity suggesting that pre-existing IgG directed against seasonal Coronavirus spike proteins are unlikely to modify interaction with SARS-CoV-2 although other cross reactive immunological mechanisms (eg T cells) cannot be ruled out and may explain the varied clinical response following exposure to SARS-CoV-2 (14). This pseudo-neutralisation assay has been shown to correlate well with neutralisation assays using live SARS-CoV-2 (MSD, personal communication). While plaque reduction neutralisation assays are currently standard for determining host antibody induced viral inhibition, they must be performed in a biosafety level 3 laboratory which limits their widespread use.

In summary, the MSD multiplexed coronavirus panel assay evaluated in this study is highly reproducible, specific and sensitive for the detection of anti-SARS-CoV-2 antibody over 14 days since the onset of COVID-19 symptoms. The assay can be adapted to measure antibody function which corelated well with spike protein antibody concentration.

## Acknowledgements

The study team would like to thank Meso Scale Discovery for the donation of the plates and reagents that allowed us to complete the work, the COSTARS study team at Great Ormond Street Children’s Hospital, staff in the Great Ormond Street Children’s Hospital Clinical Immunology Laboratory for additional support and the NIHR UCL Great Ormond Street Biomedical Research Centre for underpinning infrastructure support that facilities translation research at GOSH.

## Supplementary Tables

**Table S1:** Intra and inter-assay variability. Within plate (intra) and between plate (inter) assay repeatability was assessed by running four samples (1-4) of varying antibody levels in four replicates on the same plate and across 4 different runs on different days

| **Antigen** | **Control serum** | **Average conc. (AU)** | **Average intra-assay %CV** | **Average inter-run %CV** |
| --- | --- | --- | --- | --- |
| CoV-2 S | 1 | 2063.8 | 3.5% | 1.5% |
|  | 2 | 2579.8 | 7.2% | 1.8% |
|  | 3 | <LLOD | NA | NA |
|  | 4 | 1282.3 | 5.7% | 8.9% |
| CoV-2 RBD | 1 | 1811.0 | 4.8% | 2.1% |
|  | 2 | 2290.2 | 6.5% | 2.2% |
|  | 3 | 144.7 | 8.3% | NA |
|  | 4 | 2301.3 | 3.4% | 7.6% |
| CoV-2 N | 1 | 3380.2 | 10.9% | 0.4% |
|  | 2 | 5934.2 | 2.4% | 6.3% |
|  | 3 | <LLOD | 6.9% | 2.1% |
|  | 4 | 3557.3 | 7.5% | 7.6% |

---

**Supplementary Figure 1:**
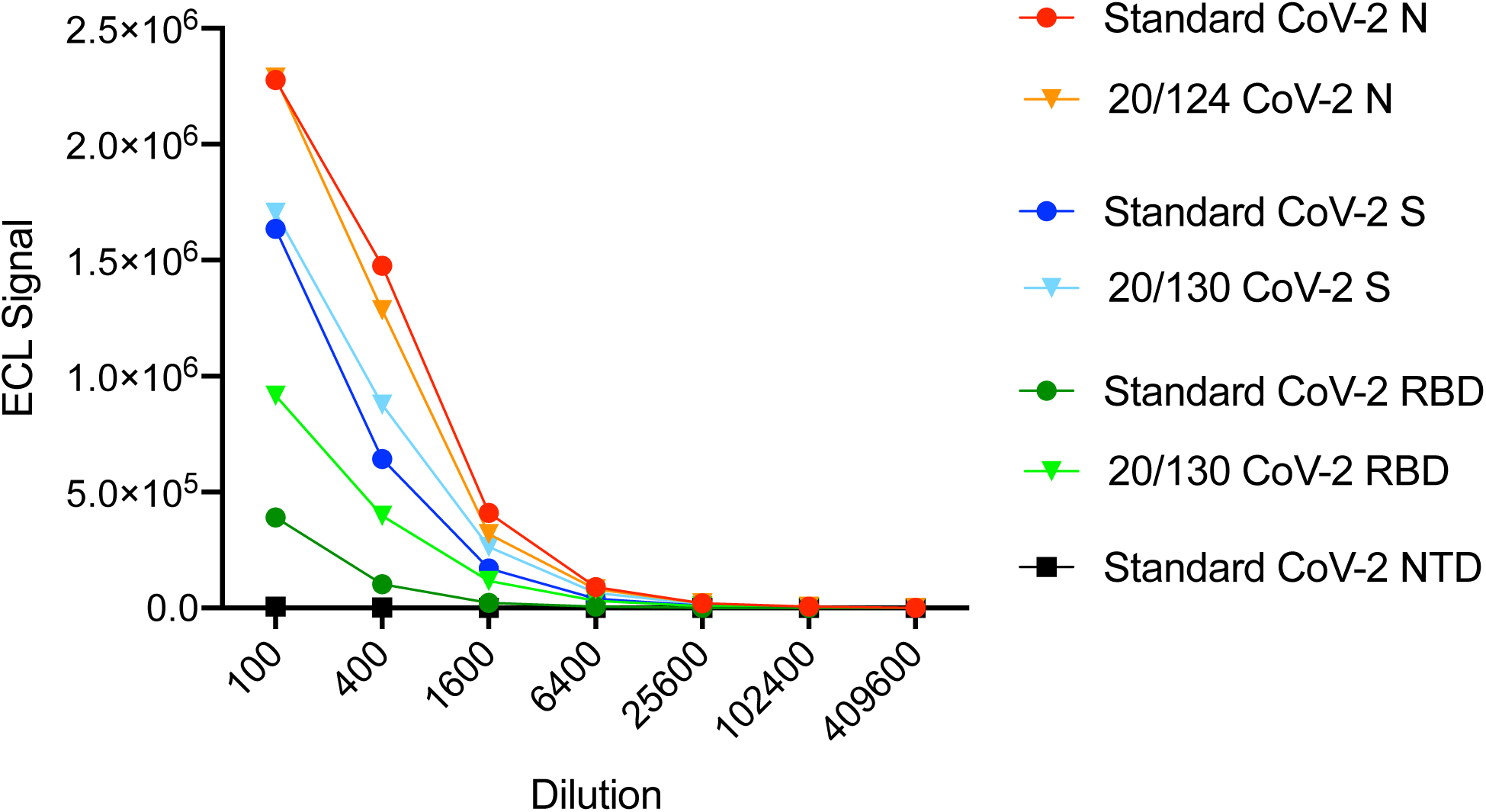
Assignment of standard values to internal standard serum and standard curves for each antigen. Graph shows ECL signal obtained from a serial dilution series (1 in 100, then 1 in 4 serial dilution) of standard serum and NIBSC control sera 20/130 and 10/124. NIBSC control serum 20/130 was used to assign values to standard serum for SARS-CoV-2 spike (S) and receptor binding domain (RBD) and NIBSC control serum 20/124 was used to assign a value to SARS-CoV-2 nucleocapsid (N). No endpoint titre corresponding to NTD antigen was available for standard serum assignment.

**Supplementary Figure 2:**
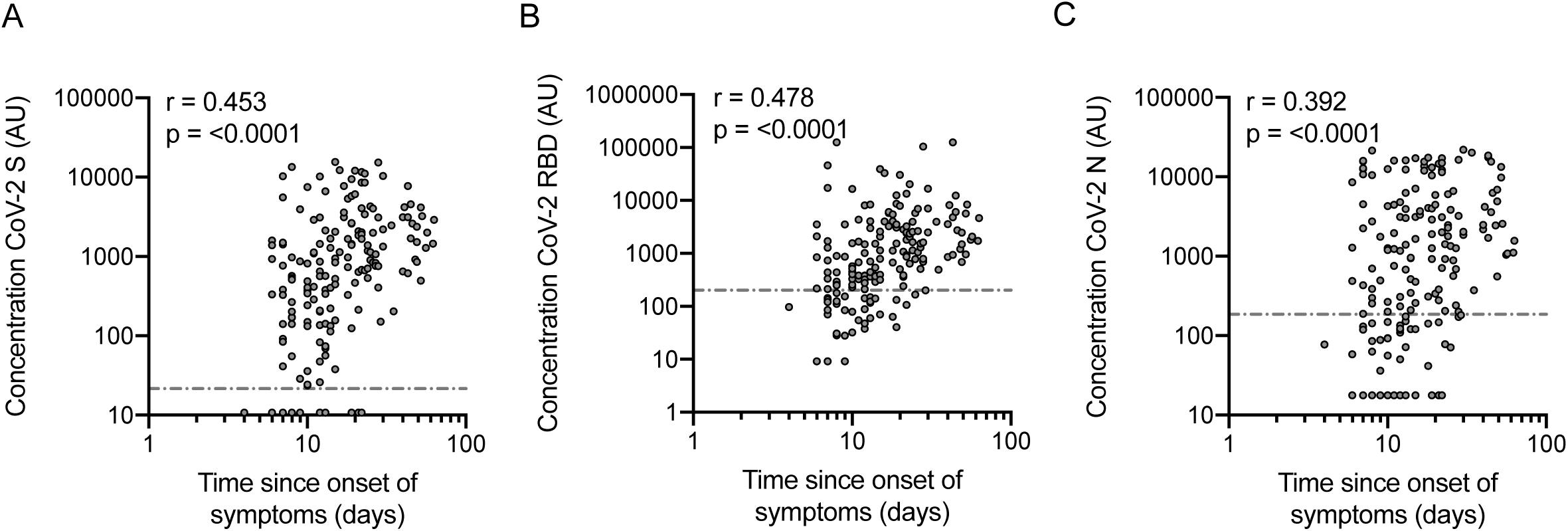
Relationship with time since onset of symptoms. Graphs show the relationship between antibody concentration against against (a) spike (S), (b) receptor binding domain (RBD) and (c) nucleocapsid (N) for all samples with known and verified time since onset of symptoms to sampling (n=176). Correlation analysis was performed using Spearman correlation. P values of <0.05 were considered as significant.

**Supplementary Figure 3:**
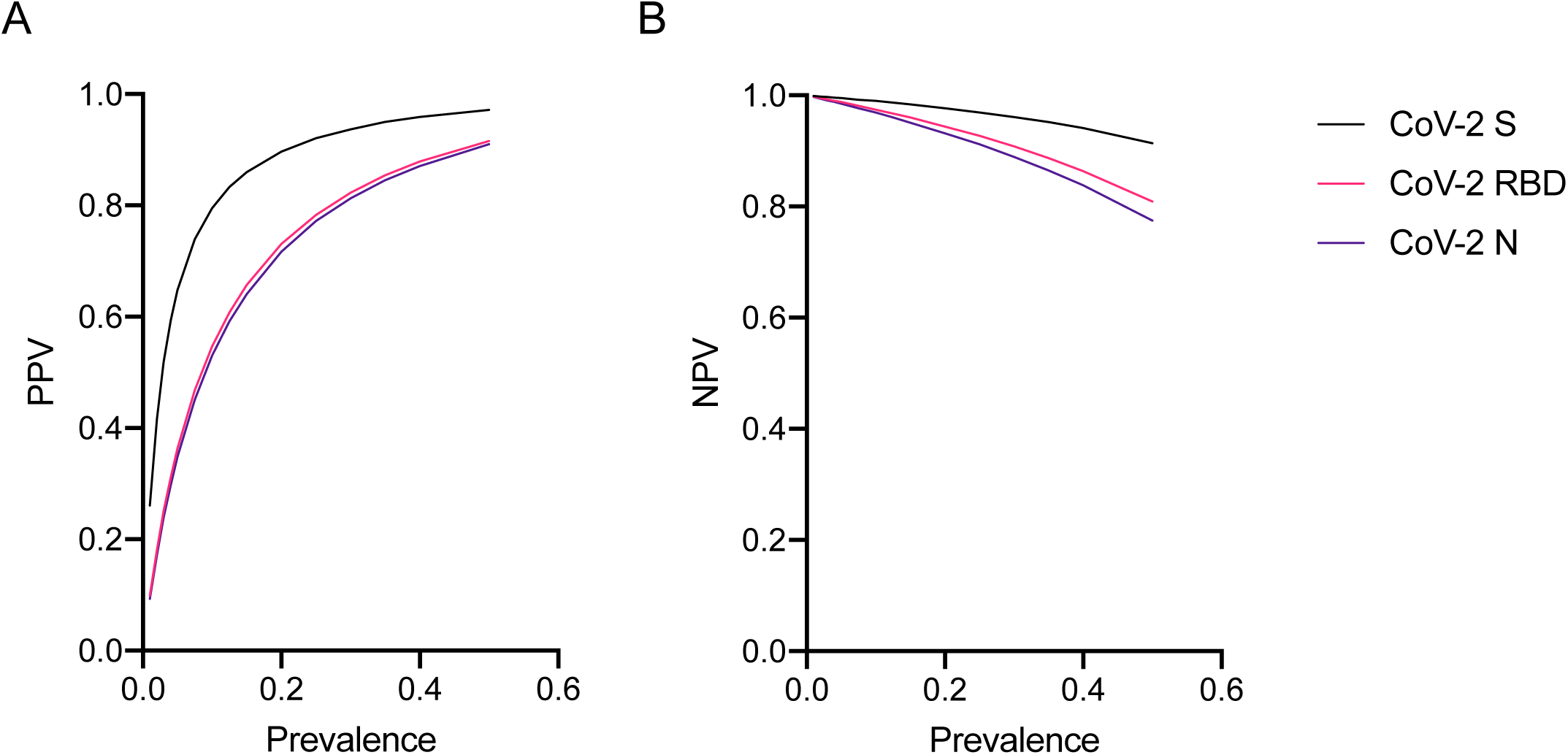
Positive and negative predictive values (PPV and NPV). Graphs show the positive (PPV) and negative (NPV) predictive values for each antigen at a range of prevalence estimates between 0.01 and 0.5 based on fixed specificity and sensitivity values calculated for the whole COVID-19 and control groups (97.4%, 92.3% and 92.8% specificity; 90.8%, 78.1% and 73.0% specificity for S, RBD and N respectively).

